# Rotavirus NSP1 subverts the antiviral OAS-RNase L pathway by inducing RNase L degradation

**DOI:** 10.1101/2022.08.24.505191

**Authors:** Jin Dai, Guanghui Yi, Asha A. Philip, John T. Patton

**Affiliations:** Department of Biology, Indiana University, Bloomington, IN 47405, USA

**Keywords:** rotavirus, interferon, OAS-RNase L, nonstructural protein 1, phosphodiesterase

## Abstract

The interferon (IFN)-inducible 2′,5′-oligoadenylate synthetase (OAS) - RNase L pathway plays a critical role in antiviral immunity. Group A rotaviruses, including the simian SA11 strain, inhibit this pathway through two activities: an E3-ligase related activity of NSP1 that degrades proteins necessary for IFN signaling, and a phosphodiesterase (PDE) activity of VP3 that hydrolyzes the RNase L-activator 2′,5′-oligoadenylate. Unexpectedly, we found that a recombinant (r) SA11 double-mutant virus deficient in both activities (rSA11-VP3H797R-NSP1ΔC17) retained the ability to prevent RNase L activation. Mass spectrometry led to the discovery that NSP1 interacts with RNase L in rSA11-infected MA104 cells. This interaction was confirmed through co-pulldown assay of cells transiently expressing NSP1 and RNase L. Immunoblot analysis showed that infection with wild type rSA11 virus, rSA11-VP3H797R-NSP1ΔC17 double-mutant virus, or single mutant forms of the latter virus, all resulted in the depletion of endogenous RNase L. The loss of RNase L was reversed by addition of the neddylation inhibitor MLN4924, but not the proteasome inhibitor MG132. Analysis of additional mutant forms of rSA11 showed that RNase L degradation no longer occurred when either the N-terminal RING domain of NSP1 was mutated or the C-terminal 98 amino acids of NSP1 were deleted. The C-terminal RNase L degradation domain is positioned upstream and is functionally independent of the NSP1 domain necessary for inhibiting IFN expression. Our studies reveal a new role for NSP1 and its E3-ligase related activity as an antagonist of RNase L and uncover a novel virus-mediated strategy of inhibiting the OAS-RNase L pathway.

**IMPORTANCE:** For productive infection, rotavirus and other RNA viruses must suppress interferon (IFN) signaling and the expression of IFN-stimulated antiviral gene products. Particularly important is inhibiting the interferon (IFN)-inducible 2′,5′-oligoadenylate synthetase (OAS) - RNase L pathway, as activated RNase L can direct the nonspecific degradation of viral and cellular RNAs, thereby blocking viral replication and triggering cell death pathways. In this study, we have discovered that the simian SA11 strain of rotavirus employs a novel strategy of inhibiting the OAS-RNase L pathway. This strategy is mediated by SA11 NSP1, a nonstructural protein that hijacks E3 cullin-RING ligases, causing the ubiquitination and degradation of host proteins essential for IFN induction. Our analysis shows that SA11 NSP1 also recognizes and causes the ubiquitination of RNase L, an activity resulting in depletion of endogenous RNase L. These data raise the possibility of using therapeutics targeting cellular E3 ligases to control rotavirus infections.

## INTRODUCTION

Rotavirus, a genus within the *Reoviridae* family, is the leading cause of acute gastroenteritis in young children (1). The virus has a non-enveloped icosahedral capsid that contains 11 segments of double-stranded (ds)RNA. These encode 6 structural (VP1-VP4, VP6-VP7) and 6 non-structural (NSP1-NSP6) proteins. During rotavirus entry, the capsid is converted into a subviral particle with associated transcriptase activity. Enzymes in the core of the subviral particle - the RNA-dependent RNA polymerase VP1 and the RNA capping enzyme VP3 - produce 11 species of capped, but non-polyadenylated plus-strand (+)RNAs. The (+)RNAs not only direct viral protein synthesis but also serve as templates for dsRNA synthesis within progeny core (2, 3). Double-stranded RNA synthesis and core formation occur within large RNA-rich inclusion bodies (viroplasms) that form in the cytoplasm of infected cells. Limiting dsRNA synthesis to progeny cores within viroplasms helps the virus avoid recognition by host dsRNA-sensors of the innate immune system. However, this is not an absolute, as studies with the dsRNA-specific J-2 antibody indicate that some rotavirus dsRNA is located outside of viroplasms (4). Moreover, several cytosolic dsRNA sensors important for triggering antiviral innate responses undergo activation during rotavirus infection, including those (e.g., RIG-I, MDA-5, PKR) that can upregulate the interferon (IFN)-signaling pathway (5, 6).

IFN expression stimulates the production of hundreds of interferon-stimulated gene (ISG) products, creating an environment unfavorable for viral replication (7). Among the most potent ISGs are those encoding the oligoadenylate synthetases (OAS). Human cells express four different isotypes of OAS [OAS1-3 and OAS-like (OASL) with OAS3 recognized as the form primarily exerting antiviral activity (8). Binding of dsRNA causes OAS1, 2 and 3 to become catalytically active, resulting in the synthesis of 2′-5′ oligoadenylate (2-5A). In the presence of 2-5A, RNase L forms dimers capable of degrading viral and cellular single-stranded RNA. The RNase L degradation products can amplify antiviral responses through recognition by cytosolic RNA sensors (e.g., RIG-I, MDA5) and activation of the apoptosis pathway (9, 10).

Group A rotaviruses (RVA), including simian SA11, can antagonize the OAS-RNase L pathway through two mechanisms, one involving the capping enzyme VP3 and the other the IFN-antagonist NSP1. VP3 hydrolyzes the OAS product 2-5A through a domain located at its C-terminus with 2’,5’ phosphodiesterase (PDE) activity (11). Site-directed mutagenesis and enzymatic analysis have indicated that VP3 PDE activity operates through two catalytic histidine residues (H718 and H797) (12, 13).

NSP1 inhibits IFN signaling and the expression of ISG products by hijacking cullin RING E3 ligases (CRLs). NSP1 recruits cellular proteins important for IFN expression to the CRL, leading to their ubiquitination and proteasomal degradation (14, 15). For example, SA11 NSP1 binds and induces the degradation of IFN regulatory factors (IRF3/5/7) (16–18). In contrast, NSP1 proteins of most human rotaviruses bind and induce the degradation of the NF-κB activator, β-transducin repeat-containing protein (β-TrCP) (19–21). NSP1 proteins may target yet an additional number of proteins for degradation, including TNF receptor-associated factor 2 (TRAF2), RIG-I, MAVS, p53 and poly(A) specific RNase subunit (Pan3) (22, 23). The NSP1 sequence is highly divergent among different rotavirus strains, a feature consistent with the variability noted in the types of cellular proteins targeted by NSP1. However, all NSP1 proteins share a conserved N-terminal RING domain, a feature essential for NSP1-mediated degradation of cellular targets (24, 25).

In this study, we have dissected mechanisms used by rotavirus to subvert the OAS-RNase L pathway. Through analysis of recombinant (r) SA11 rotaviruses encoding VP3 with a defective PDE domain and NSP1 unable to inhibit IFN signaling, we discovered an alternative mechanism used by SA11 rotavirus to inhibit the OAS-RNase L pathway. This mechanism is mediated by NSP1, which binds and causes the degradation of RNase L through a ubiquitination-dependent but proteasome-independent process. Although other viruses are known to produce proteins that inhibit the OAS-RNase L pathway, this is the first example of a virus blocking the pathway through degradation of its RNase L component.

## RESULTS

### Contribution of VP3 PDE and NSP1 activities to rotavirus replication

To investigate the importance of VP3 PDE and NSP1 activities on rotavirus biology, we used reverse genetics to generate rSA11 rotaviruses expressing inactive forms of VP3 PDE and NSP1. A virus expressing inactive VP3 PDE was produced by mutating one of the catalytic histidine residues in the PDE domain to arginine (rSA11-VP3H797R) (12) (Fig. 1A). A virus defective in suppressing IFN expression was generated by modifying the NSP1 open reading frame such that the protein product contained a C-terminal 17 amino acid truncation (rSA11-NSP1ΔC17) (17) (Fig. 1A). Given that VP3 PDE and NSP1 may work together to suppress immune responses, we also generated a recombinant virus expressing both inactive VP3 PDE and NSP1 (rSA11-VP3H797R-NSP1ΔC17). Recovery of mutant viruses was verified by RNA gel electrophoresis (Fig. 1B) and cDNA sequencing. Immunoblot analysis with NSP1 antibody confirmed that rSA11-NSP1ΔC17 and rSA11-VP3H797R-NSP1ΔC17 expressed a truncated form of NSP1 (Fig. 1C).

**Figure 1.**
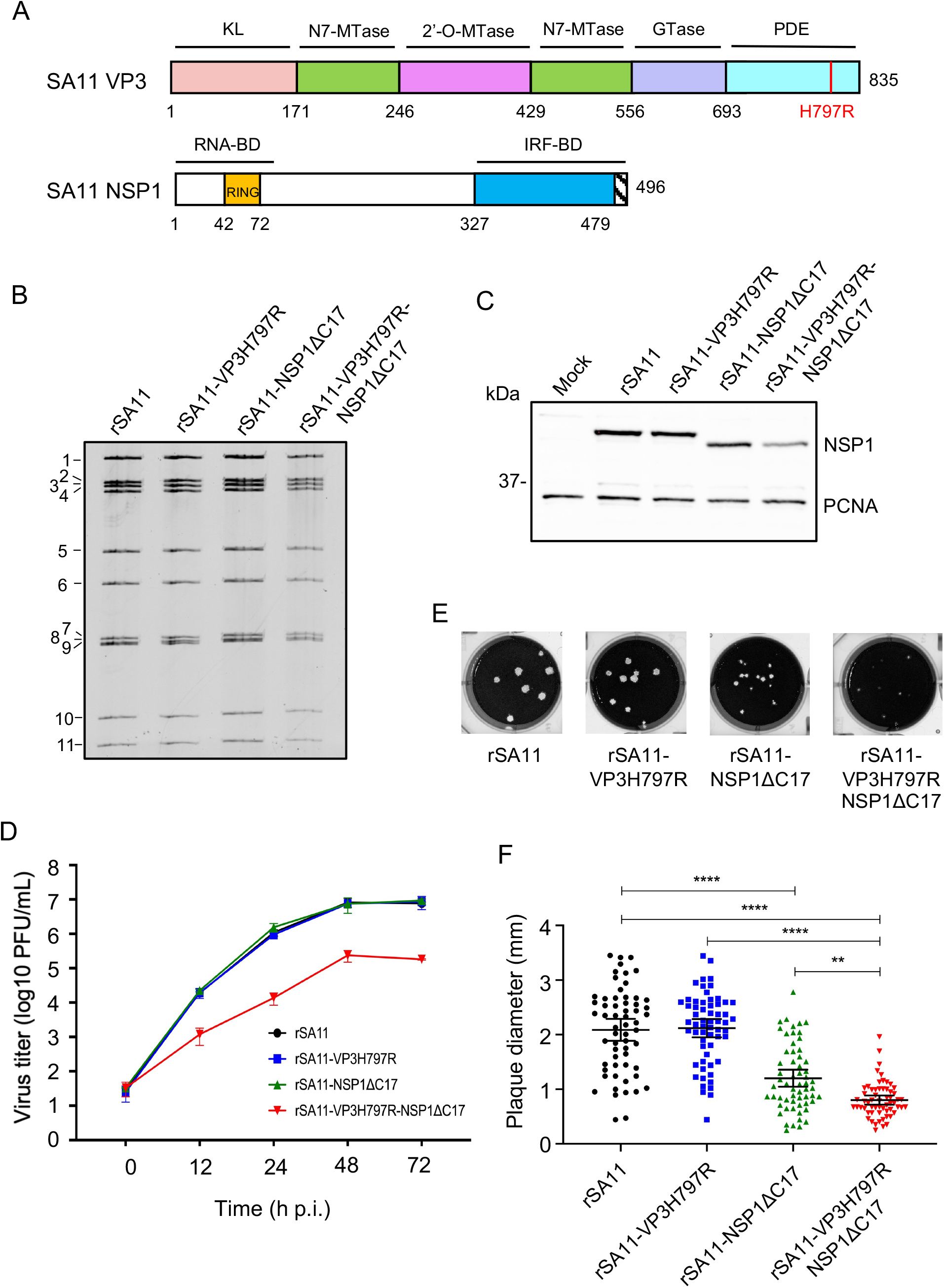
Importance of VP3 PDE and NSP1 to efficient rotavirus replication. (A) Domains of SA11 VP3 and NSP1. VP3: pink, kinase-like (KL); green, N7-methyltransferase (N7-MTase); magenta, 2’-O-methyltransferase (2’-O-MTase); lavender, guanylyltransferase (GTase); cyan, 2’-5’ phosphodiesterase (PDE). NSP1: yellow, RING domain; blue, IRF-binding domain (BD). Mutations in VP3 (H797R) and NSP1 (deletion of 479-496, ΔC17) are indicated. (B) Electrophoretic profiles of genomic dsRNA from wildtype (rSA11) and mutant rotaviruses. (C) Immunoblot analysis of NSP1 and PCNA in mock- or virus-infected MA104 cells at 10 hpi. (D) Multistep growth curves of wildtype and mutant rotaviruses (MOI = 0.01) in MA104 cells. Virus titers were determined by plaque assay from two independent experiments. (E) Plaques formed by wildtype and mutant rotaviruses on MA104 cells. (F) Plaques sizes formed by wildtype and mutant rotaviruses. Total of 60 plaques were measured by ImageJ software from three independent experiments. Black lines represent the mean ± 95% confidence intervals. Significance values were calculated using a oneway ANOVA test. **, *P* ≤ 0.01; ****, *P* ≤ 0.0001.

To explore the impact of VP3 PDE and NSP1 mutations on rotavirus growth, we compared multistep growth curves produced for VP3H797R, NSP1ΔC17, and VP3H797R-NSP1ΔC17 viruses using the simian MA104 cells (Fig. 1D). The analysis showed that the VP3H797R and NSP1ΔC17 single mutant viruses replicated at rates similar to wild type (WT) rSA11 virus. However, the VP3H797R-NSP1ΔC17 double mutant virus grew less efficiently in comparison to WT or the VP3H797R or NSP1ΔC17 single mutant viruses (Fig. 1D). We also compared the plaques formed by these viruses on MA104 cells. Consistent with previous studies (17), the NSP1ΔC17 virus produced plaques smaller than those of WT virus. In contrast, the VP3H797R virus produced plaques similar in size to WT virus. Interestingly, plaques formed by the VP3H797R-NSP1ΔC17 double mutant virus were even smaller than those of the NSP1ΔC17 virus, suggesting that VP3 PDE contributes to efficient virus replication or spread through a mechanism that is not dependent on functional NSP1 (Fig. 1E,F). Through immunoblot analysis of ISGs (IFIT1 and viperin), we determined that the VP3H797R-NSP1ΔC17 double mutant virus stimulated higher levels of ISG expression than either the VP3H797R or the NSP1ΔC17 single mutant viruses (Fig. S1). This indicates that VP3 PDE plays a valuable role in suppressing IFN/ISG pathways that extends beyond the capabilities of NSP1 alone.

### Rotavirus inhibition of RNase L activation does not require VP3 PDE or NSP1 suppression of IFN signaling

To evaluate how the VP3H797R, NSP1ΔC17, and VP3H797R-NSP1ΔC17 mutations affected RNase L activation during rotavirus infection, the integrity of rRNAs in MA104 cells infected with these viruses was examined using an RNA TapeStation assay. In agreement with previous studies, 18S and 28S rRNA remained intact in cells infected with WT virus suggesting that rotaviruses can evade the OAS-RNase L pathway (26, 27). Surprisingly, no rRNA degradation was observed in cells infected with VP3H797R or NSP1ΔC17 single mutant viruses. Even infection with the VP3H797R-NSP1ΔC17 double mutant virus did not induce obvious rRNA degradation (Fig. 2A). To exclude the possibility that the OAS-RNase L pathway might be defective in MA104 cells, we examined rRNA integrity in these cells following transfection with purified reovirus dsRNA. Consistent with a functional OAS-RNase L pathway, degradation of rRNA was apparent in the dsRNA-transfected MA104 cells (Fig. 2B). To test whether rotavirus prevents RNase L activation even in the presence of a dsRNA stimulus, we infected MA104 cells with WT, VP3H797R, NSP1ΔC17, and VP3H797R-NSP1ΔC17 viruses, then transfected these cells with viral dsRNA. The results showed that despite transfection with dsRNA, rRNA was not degraded in cells infected with any of these viruses, including the VP3H797R-NSP1ΔC17 double mutant that lacks a functional VP3 PDE and cannot suppress IFN expression (Fig. 2B). These observations suggest that, apart from its ability to degrade 2-5A and suppress IFN expression, rotavirus uses other mechanism(s) to protect itself from RNase L.

**Figure 2.**
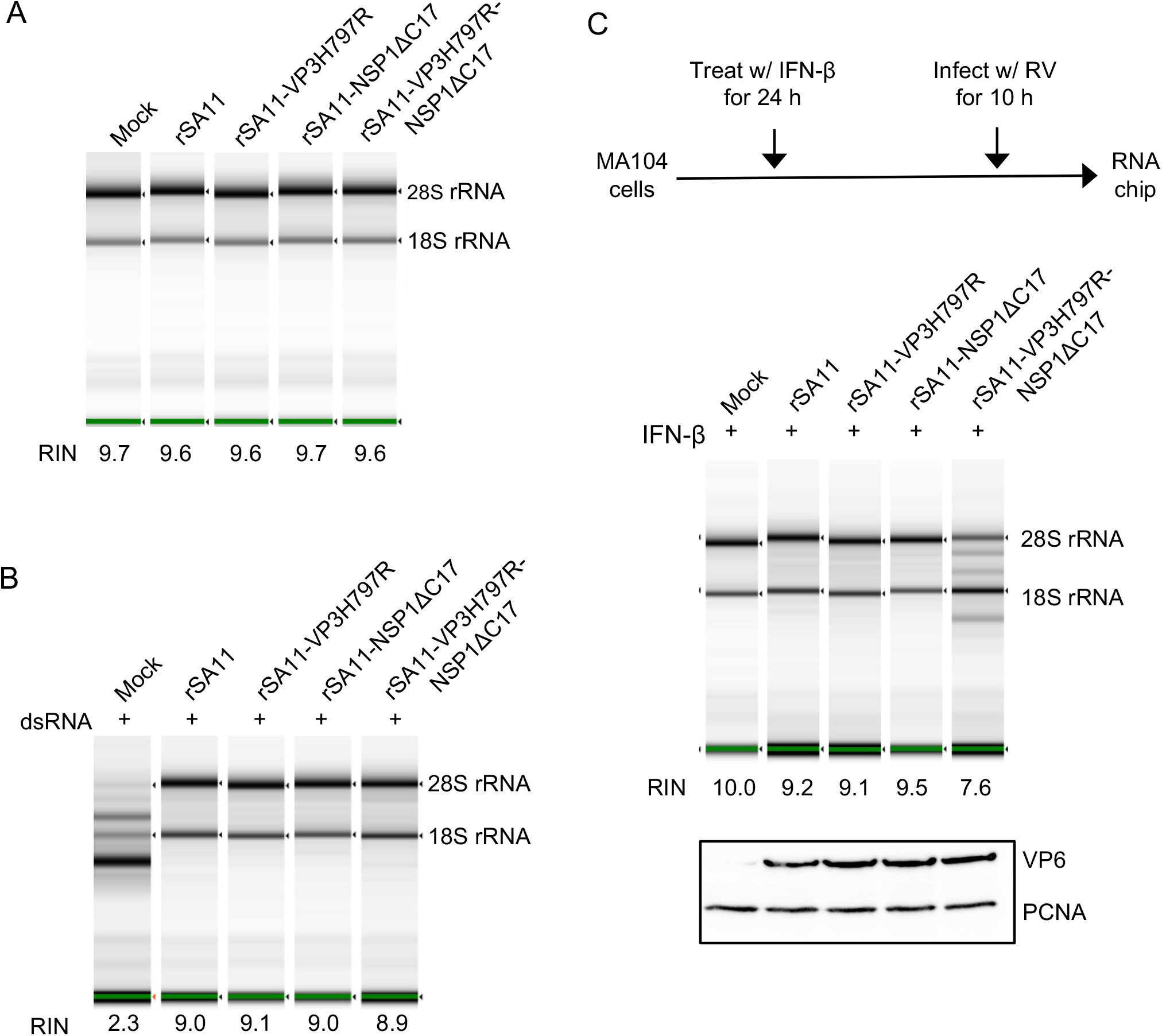
Effect of VP3 PDE and NSP1 on ribosomal (r) RNA degradation in rotavirus-infected cells. (A) MA104 cells were mock-infected or infected with the indicated rotavirus strains. Cellular RNA was recovered from cells harvested at 10 h p.i. and analyzed by automated electrophoresis using an Agilent TapeStation 2200 system. RNA integrity number (RIN) was calculated using TapeStation software. (B) MA104 cells were infected with the indicated rotavirus strains and transfected with 500 ng of dsRNA at 5 h p.i. RNA was extracted at 10 h p.i. and analyzed using the TapeStation. (C) MA104 cells were pretreated with rhesus monkey IFN-β at 24 h prior to infection with indicated rotavirus strains (MOI = 6). RNA recovered from cells at 10 h p.i. analyzed using the TapeStation. Immunoblot assay was used to detect VP6 and PCNA in cells harvested at 10 h p.i.

During natural infection, paracrine IFN signaling can trigger the expression of antiviral ISGs, such as OAS, in neighboring uninfected cells (28). To determine if such paracrine IFN signaling could promote RNase L activation in a way that could not be subverted by rotavirus, MA104 cells were treated with IFN-β and then infected with WT or mutant viruses. Ten hours later, rRNA integrity in those cells was examined by RNA TapeStation assay. The results showed that infection with WT virus and VP3H797R and NSP1ΔC17 single mutant viruses did not activate RNase L despite pre-treatment of the cells with IFN-β. In contrast, rRNA were degraded in cells infected with the VP3H797R-NSP1ΔC17 double mutant virus, indicating that rotavirus dsRNA can activate the OAS-RNase L pathway, but through a process requiring prior exposure to type I interferon (Fig. 2C). These results support the idea that rotavirus has multiple mechanisms of preventing RNase L activation.

One explanation for the failure of the VP3H797R-NSP1ΔC17 double mutant virus to prevent activation of the OAS-RNase L pathway was that the virus expressed insufficient levels of viral proteins in IFN-treated cells. To examine this possibility, we compared levels of the rotavirus VP6 protein produced in cells infected with WT and mutant viruses by immunoblot assay. The results indicated that these viruses expressed comparable levels of VP6 (Fig. 2C), suggesting that RNase L activation by the VP3H797R-NSP1ΔC17 double mutant virus did not result from reduced viral protein expression.

### Degradation of RNase L during rotavirus infection

The ability of the VP3H797R-NSP1ΔC17 double mutant virus to protect rRNA from degradation was equivalent to that seen with WT virus (Fig. 2B). This observation led us to ask whether rotavirus employs a third mechanism to inhibit the OAS-RNase L pathway. To address this question, we infected HT-29 cells with recombinant rSA11 viruses expressing 3×FLAG-tagged VP3 (rSA11-VP3-3×FLAG) or 3xFLAG-tagged NSP1 (rSA11-NSP1-3×FLAG-UnaG). Following lysis, immunoprecipitates containing 3xFLAG-tagged VP3 or NSP1 were recovered using anti-FLAG beads. The immunoprecipitates were analyzed by mass spectrometry to define the interactomes of VP3 and NSP1. Notably, RNase L was detected in the immunoprecipitate recovered for 3×FLAG-tagged NSP1, but not detected in the immunoprecipitate recovered for 3×FLAG tagged VP3. Given that NSP1 is known to induce the degradation of multiple host proteins, including IRF3 and β-TrCP (22), we examined the possibility that NSP1 might also target RNase L for degradation. To address this question, MA104 cells were infected with WT virus and with the VP3H797R, NSP1ΔC17, and VP3H797R-NSP1ΔC17 mutant viruses. Lysates prepared from the cells at 8 h p.i. were then analyzed for endogenous RNase L by immunoblot assay. The results showed that all the viruses caused depletion of endogenous RNase L, even the double mutant encoding both defective forms of VP3 PDE and NSP1 (Fig. 3A).

**Figure 3.**
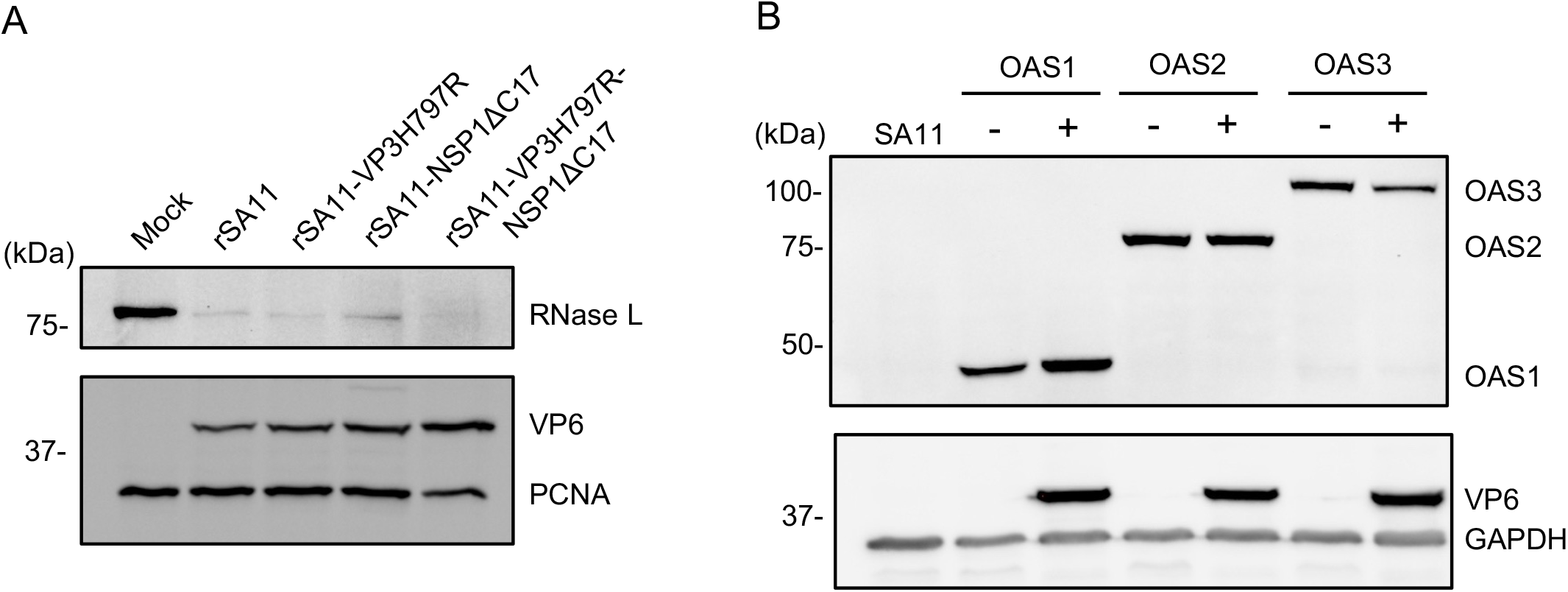
Depletion of RNase L during rotavirus infection. (A) MA104 cells were infected with the indicated strains of rotavirus (MOI = 6) and harvested at 8 h p.i. Immunoblot assay was used to detect RNase L, PCNA and rotavirus VP6 in cell lysates. (B) HT-29 cells were transfected with plasmids encoding the indicated OAS proteins and, 24 h later, infected with rSA11 (MOI = 5). Immunoblot assay was used to detect OAS1-3, VP6 and GAPDH in the cell lysates. Blots are representative of the results of three independent experiments. Sizes of protein markers (kDA) are indicated.

To determine whether rotavirus infection caused depletion of other proteins necessary for activation of the OAS-RNase L pathway, we transfected the human colon HT-29 cells with plasmids expressing FLAG-tagged human OAS1, 2, or 3. Twenty-four hours later, the cells were infected with WT rSA11 viruses. Immunoblot analysis of lysates prepared from these cells indicated that rotavirus infection had no impact on levels of transiently expressed OAS1, 2, or 3 (Fig. 3B).

To further investigate the role of NSP1 in the depletion of RNase L, we infected HT-29 cells with a recombinant SA11 rotavirus expressing only the first 35 residues of the 496-aa NSP1 protein (rSA11-NSP1-35TAA). Unlike its WT counterpart, this NSP1 deficient virus did not cause significant RNase L depletion, indicating that NSP1 is responsible for loss of the RNase L protein (Fig. 4A). To validate this conclusion, we transiently expressed FLAG-tagged human RNase L and Halo-tagged SA11 NSP1 together in 293T cells. Twenty-four hours post-transfection, the cells were harvested and analyzed by immunoblot assay for FLAG-tagged RNase L. The results showed that FLAG-tagged RNase L was lost only when co-expressed with Halo-tagged NSP1 (Fig. 4B).

**Figure 4.**
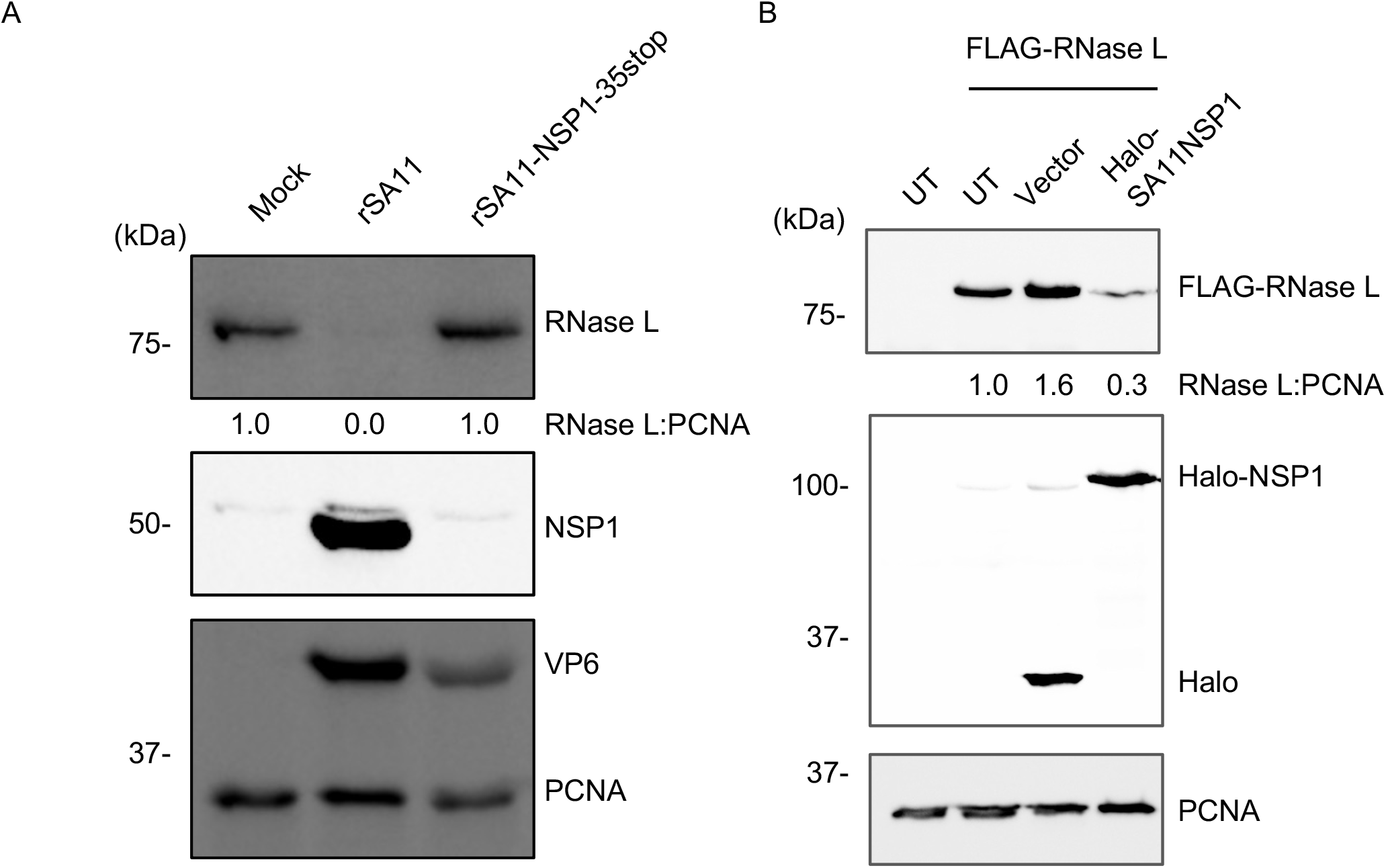
SA11 NSP1 causes reduced RNase L levels. (A) HT-29 cells were mock infected or infected with the indicated rotaviruses (MOI = 5). Proteins in cell lysates were detected by immunoblot assay using antibodies specific to RNase L, NSP1, VP6, and PCNA. (B) HEK 293T cells were cotransfected with a plasmid expressing FLAG-tagged human RNase L and a plasmid encoding Halo-tagged SA11 NSP1 or an empty pHTN vector. Proteins in cell lysates were collected at 24 h p.t. and analyzed by immunoblot assay using antibodies for HaloTag, FLAG, and PCNA. Blots are representative of the results of three independent experiments.

### Neddylation inhibitor MLN4924 partially rescues RNase L

Previous studies have shown that the NSP1 can hijack CRLs, leading to the ubiquitination and proteasomal degradation of host proteins important for antiviral responses (e.g., β-TrCP, IRF3, IRF7) (18-21). To further dissect the molecular mechanisms underlying NSP1-mediated RNase L degradation, we tested whether inhibitors targeting the functions of CRLs (MLN4924), proteasomes (MG-132) and lysosomes (chloroquine) could reverse the depletion of RNase L in rSA11-infected HT-29 cells. Immunoblot assay showed that only MLN4924 was able to rescue RNase L in HT-29 cells, suggesting the NSP1 induces RNase L depletion in a ubiquitination-dependent manner. Surprisingly, MG-132, chloroquine and even the combination of these two inhibitors did not rescue RNase L, suggesting rSA11 may use a novel mechanism of targeting RNase L for degradation (Fig. 5A). To confirm these results, 293T cells were treated with MLN4924 and then transfected with plasmids expressing FLAG-tagged human RNase L and Halo-tagged SA11 NSP1. Consistent with the results above, MLN4924 treatment prevented transiently expressed NSP1 from inducing depletion of RNase L (Fig. 5B).

**Figure 5.**
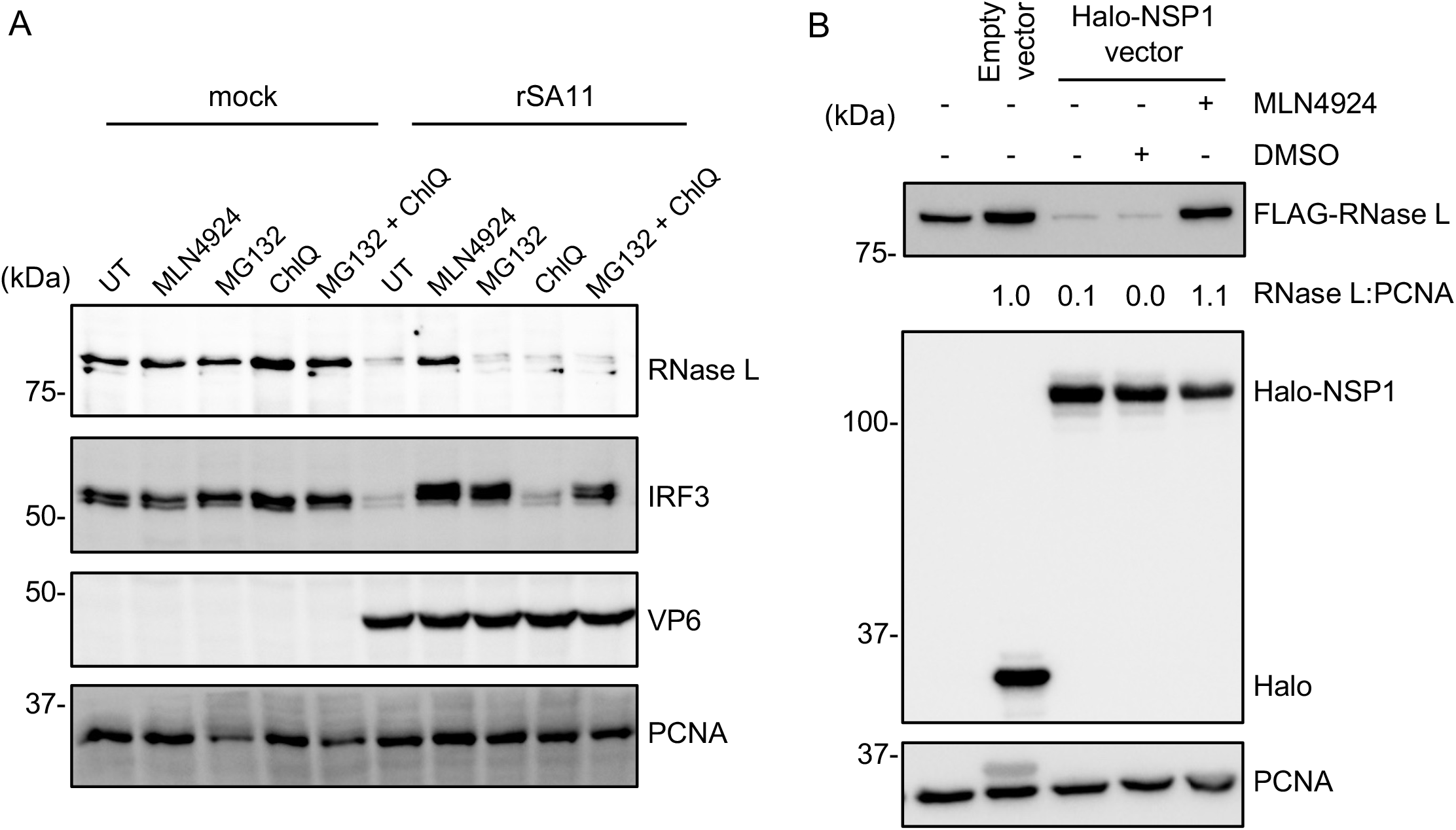
Neddylation inhibitor MLN4924 partially rescues RNase L. (A) HT-29 cells were treated with the indicated inhibitors and infected with rSA11 (MOI = 5). Cells were harvested at 8 h p.t. and analyzed by immunoblot assay for RNase L, IRF3, VP6, and PCNA. UT, untreated; ChlQ, chloroquine. (B) HEK 293T cells were treated with MLN4924 or DMSO. At 1 h post-treatment, cells were co-transfected with a plasmid encoding FLAG-tagged human RNase L and a plasmid expressing Halo-tagged SA11 NSP1 or an empty pHTN vector. Cells were harvested at 24 h p.t. and analyzed by immunoblot assay using antibodies specific for HaloTag, FLAG tag, and PCNA. Blots are representative of the results of three independent experiments.

### NSP1 interacts with RNase L

To confirm the physical interaction between NSP1 and RNase L, Halo-tagged SA11 NSP1 and FLAG-tagged human RNase L were transiently expressed in 293T cells pretreated with MLN4924. At 24 h post-transfection, the cells were lysed and incubated with anti-FLAG beads. The beads were recovered, and a portion treated with RNase A and RNase T1 to remove proteins bound through RNA-bridging interactions. Immunoblot assay using anti-Halo tag antibody showed that NSP1 was present in FLAG-tagged RNase-L immunoprecipitates irrespective of the RNase treatment (Fig. 6A). Reciprocally, we performed a pull-down assay using HaloLink resin to recover proteins associated with Halo-tagged NSP1. The results showed that RNase L was present in the Halo-tagged NSP1 immunoprecipitates even after RNase treatment (Fig. 6B).

**Figure 6.**
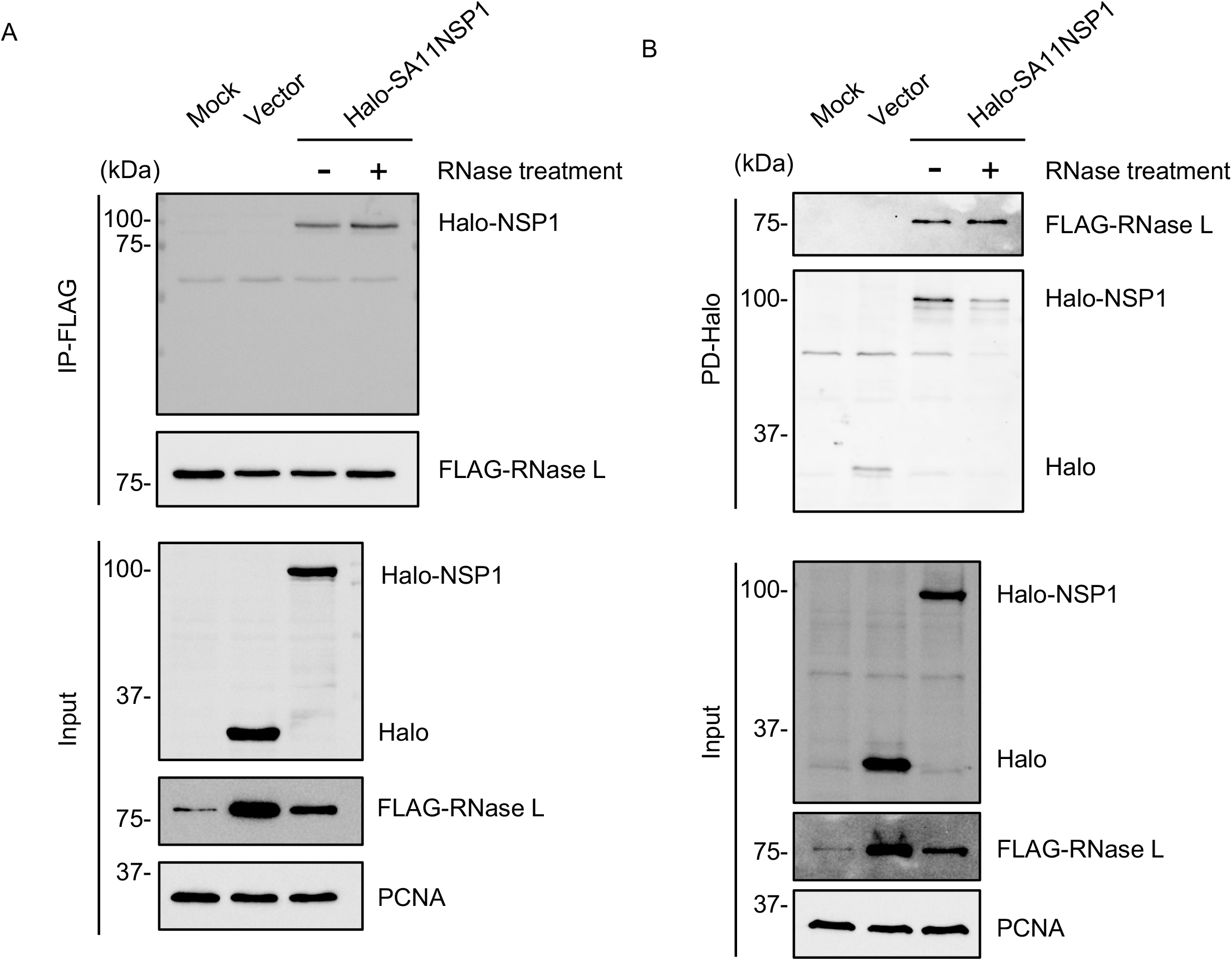
NSP1 interacts with RNase L. (A) HEK 293T cells were treated with MLN4924, and 1 h later, co-transfected with a plasmid expressing FLAG-tagged RNase L and a plasmid expressing Halo-tagged SA11 NSP1 or an empty vector. Lysates prepared from the cells at 24 h p.t. were incubated with magnetic beads conjugated with mouse anti-FLAG antibody to immunoprecipitate complexes containing FLAG-tagged proteins. The beads were split into two equal aliquots, with one aliquot treated with RNase A and RNase T1 cocktail. Proteins bounds to beads and input fractions were analyzed by immunoblot assay using antibodies to FLAG, HaloTag, and PCNA. (B) Reciprocal immunoprecipitation of (A) performed with anti-HaloTag resin instead of magnetic beads conjugated to anti-FLAG antibody. Blots are representative of the results of two independent experiments.

### NSP1 domains required for RNase L degradation

Current models suggest that both the C terminal substrate recognition domain and the N terminal RING domain of NSP1 are necessary for NSP1-induced degradation of IRF3 (16) and β-TrCP (21, 25). To map the regions of NSP1 necessary for RNase L degradation, plasmids were generated that expressed Halo-tagged NSP1 with C-terminal truncations of 17, 98, 243, or 461 residues or Halo-tagged NSP1 with a C42A mutation in the RING domain (Fig. 7A). 293T cells were co-transfected with plasmids encoding mutant forms of Halo-tagged NSP1 and a plasmid encoding FLAG-tagged human RNase L. The ability of the different forms of NSP1 to induce the degradation of RNase L was analyzed by immunoblot assay. Like results observed with WT or rSA11-NSP1ΔC17 infected MA104 cells (Fig. 3A), plasmids expressing full-length and ΔC17 NSP1 caused RNase L depletion (Fig. 7B). In contrast, expression of NSP1 with C-terminal truncations of 98, 243 or 461 residues did not result in the loss of RNase L, suggesting that a region in the last 98 residues is essential for the targeting of RNase L. NSP1 with a C42A RING mutation also failed to cause RNase L degradation, suggesting the RING domain also plays an essential role in targeting RNase L (Fig. 7B).

**Figure 7.**
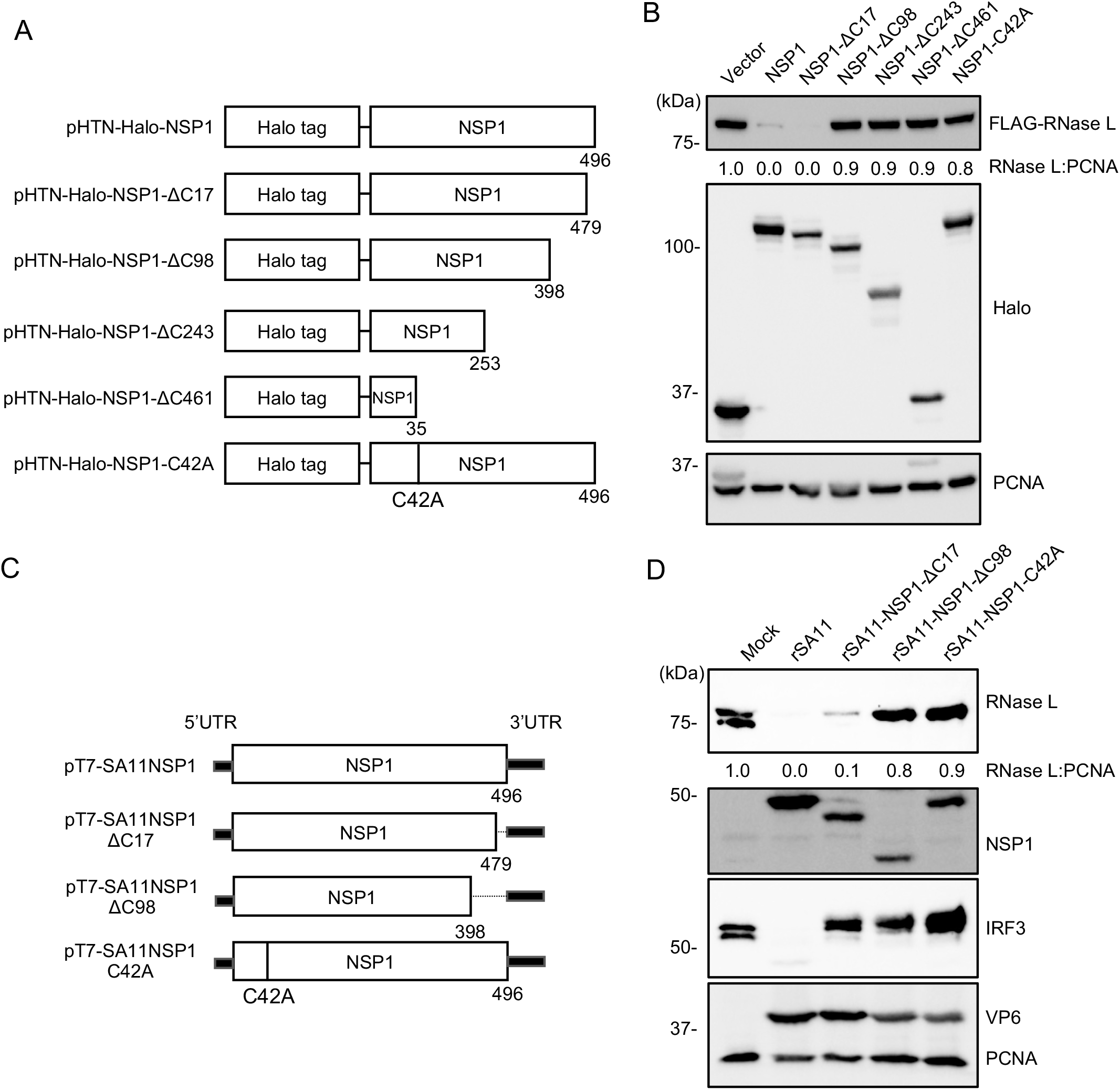
Regions of NSP1 contributing to RNase L degradation. (A) Schematic of pHTN plasmids encoding mutant Halo-tagged SA11 NSP1 proteins. (B) HEK 293T cells were cotransfected with a plasmid encoding FLAG-tagged RNase L and a plasmid encoding a Halo-tagged NSP1 protein. Cell lysates were prepared at 24 h p.t. and analyzed by immunoblot assay for HaloTag, FLAG, and PCNA. (C) Schematic of pT7 plasmids encoding wildtype or mutant NSP1, used in recovering wildtype and mutant rotaviruses. (D) HT-29 cells were infected with indicated rotavirus strains (MOI = 5). Cells were harvested at 8 h p.i. and analyzed by immunoblot assay for RNase L, NSP1, VP6, and PCNA. Blots are representative of three independent experiments.

To validate these results in the context of rotavirus infection, rSA11 viruses were generated that expressed NSP1 with a C-terminal truncation of 98 residues (rSA11-NSP1-ΔC98) or with a C42A RING mutation (rSA11-NSP1-C42A) (Fig. 7C). Analysis of HT-29 cells infected with these two mutant viruses, showed that neither caused degradation of endogenous RNase L, in contrast to WT or rSA11-NSP1ΔC17 viruses (Fig. 7D). These results suggest that the RING domain and a region close to the C-terminus of SA11 NSP1 are required for RNase L depletion. Interestingly, although the last 17 residues of SA11 NSP1 are essential for IRF3 degradation, this region is not required for RNase L degradation.

### RNase L degradation activity is not a shared feature of the group A rotaviruses

Previous studies have shown that NSP1 proteins vary in the types of cellular proteins that they target for degradation (24). To determine whether RNase L degradation is a shared feature of rotavirus NSP1 proteins, we infected HT-29 cells with four common laboratory strains (rSA11, SA11-L2, RRV and OSU) and a panel of monoreassortant viruses containing genome segment 5 (NSP1) dsRNAs derived from different rotavirus strains on the genetic background of the SA11-L2 virus (29). Immunoblot analysis showed that only the NSP1 proteins of the simian rSA11 and SA11-L2 strains possessed RNase L degradation activity. Even another simian rotavirus strain (RRV) and an SA11 reassortant strain expressing RRV NSP1 did not target RNase L (Fig. 8).

**FIG 8.**
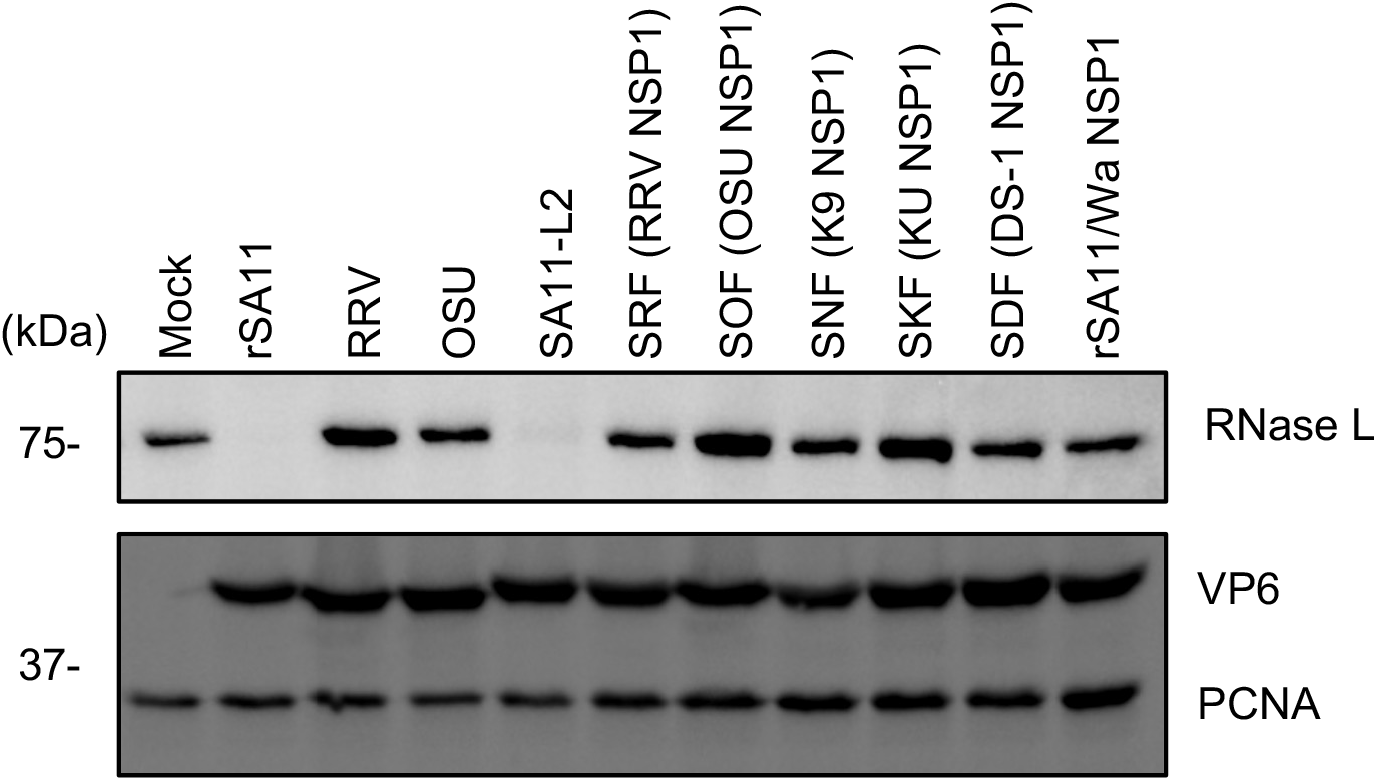
Rotavirus strain type and loss of RNase L. Human HT-29 cells were infected with the indicated rotavirus strain (MOI = 5). Cells were harvested at 8 h p.i. and analyzed by immunoblot assay with antibodies against RNase L, VP6, and PCNA. The data are representative of three independent experiments. SRF, SOF, SNF, SKF, and SDF strains are SA11 reassortants with NSP1 segments derived from RRV, OSU, K9, KU, and DS-1.

## DISCUSSION

Previous studies have shown that rotaviruses use two strategies to suppress activation of the OAS-RNase L pathway: (i) degradation of 2-5A by VP3 PDE and (ii) inhibition of OAS expression by the IFN-antagonist NSP1. Unexpectedly, we found that a double-mutant SA11 virus deficient in both activities retained the ability to prevent RNase L activation in the infected cell. Analysis of this alternative strategy indicates that it is mediated by SA11 NSP1, via a process in which NSP1 directly binds to and ultimately causes ubiquitination-dependent but proteasome-independent RNase L degradation (Fig. 9). This alternative strategy allows the SA11 virus to suppress the OAS-RNase L pathway even in infected cells containing activated OAS producing 2-5A.

**Fig. 9.**
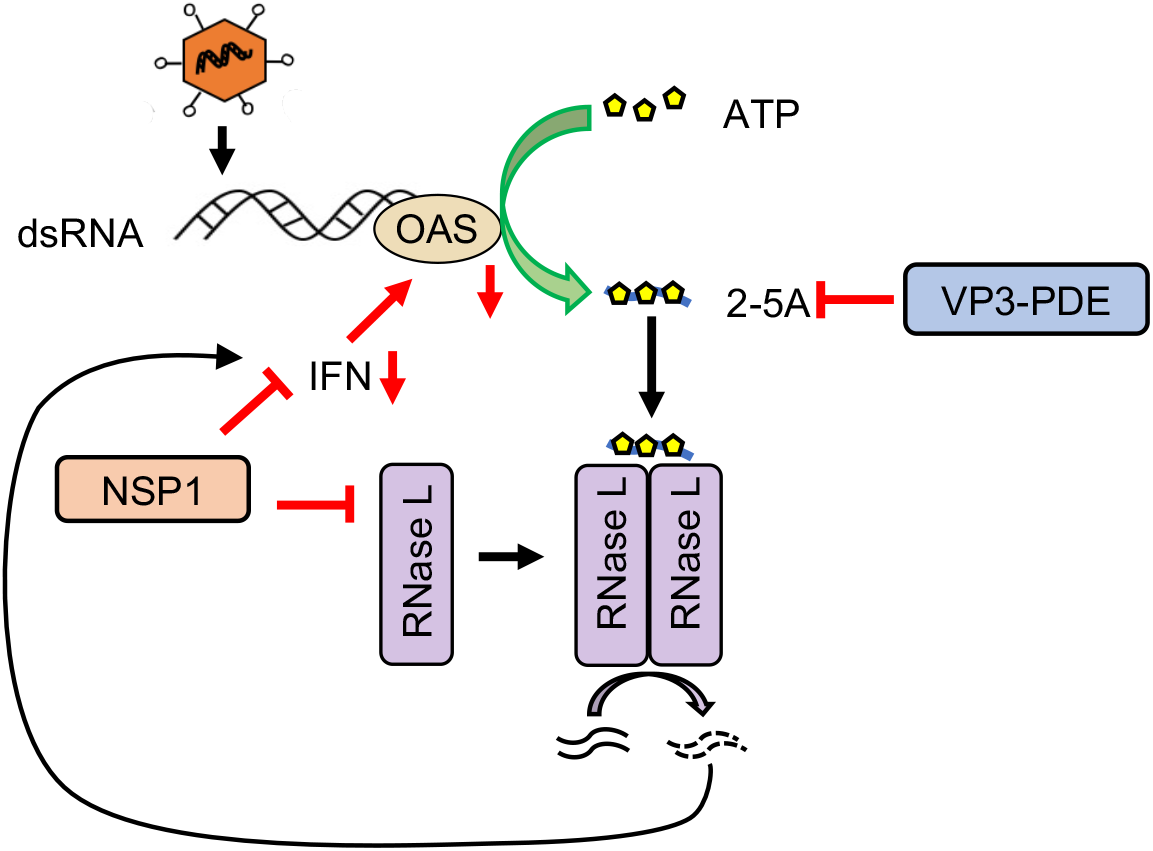
Strategies used by rotavirus SA11 to evade the OAS-RNase L pathway. The three mechanisms are degradation of 2-5A by VP3 PDE, inhibition of the IFN-signaling pathway by NSP1, and induction of RNase L degradation by NSP1.

Previous studies have indicated that by hijacking CRLs, rotavirus NSP1 induces the ubiquitination and degradation of cellular proteins critical to innate immune responses, such as IRF3/7 and β-TrCP. Features of NSP1 critical to this function are its N-terminal RING domain, which possibly plays a role in recruiting ubiquitin charged accessory proteins, and its C-terminal targeting domains, which direct interactions with cellular proteins (24). Given that RNase L degradation also requires an intact RING domain and involves a C-terminal targeting domain, NSP1 seems to employ a similar mechanism for targeting RNase L, IRF3/7, and β-TrCP. Indeed, degradation of all three is inhibited by the neddylation inhibitor MLN4924 (20), pointing to the involvement of an CRL-like activity that mediates target-protein ubiquitination. Although experiments with the inhibitor MG-132 indicate that IRF3/7 and β-TrCP undergo proteasomal degradation, this inhibitor nor an lysosome inhibitor (chloroquine) had any impact on RNase L degradation, suggesting a less typical route of degradation. Notably, the RNase L targeting domain in SA11 NSP1 is located upstream of the IRF3/IRF7 targeting domain and can function even in the absence of the IRF3/7 targeting domain. The nature of the targeting domains accumulated for any one NSP1 protein likely reflects variation in the selective pressures imposed on virus strains replicating in different host species.

The strategies taken by the various groups of rotaviruses to antagonize the OAS/RNase L pathway are markedly different. Unlike SA11 and other group A rotaviruses, group C rotaviruses do not encode VP3 PDE. Instead, one of the genome segments (NSP3) of group C viruses encodes a small extra protein with dsRNA-binding activity. This protein is known to inhibit PKR activation and may similarly act to inhibit OAS activation by sequestering viral dsRNA (30).

Although the activities used by the SA11 virus to inhibit the OAS-RNase L pathway seem rather redundant, this may be necessary for productive virus replication and spread in an intestinal environment where some cells are naïve, but others have entered into an antiviral state due to paracrine IFN signaling. Moreover, because some cells are known to produce high basal levels of OAS, even in the absence of IFN (31, 32), the ability of SA11 NSP1 to prevent IFN production may not be sufficient. The fact that SA11 has two specific-targeting mechanisms for shutting down the OAS-RNase L pathway may have resulted from their combined ability to prevent not only RNase L activation but also induction of ISGs (Fig. S1). It is also worth noting that VP3 PDE is a component of a structural protein (VP3) that is expressed at low levels in infected cells and constantly being depleted through its uptake into viroplasms and packaging into progeny cores. In contrast, albeit like VP3 expressed at low levels, the nonstructural protein NSP1 is expressed throughout the infection and accumulates outside the viroplasm.

Degradation of RNase L represents a novel, previously unrecognized, viral strategy of subverting the OAS-RNase L pathway. Other viral strategies include expression of dsRNA-binding proteins that sequester viral dsRNAs, impeding dsRNA-dependent OAS activation OAS. Examples are the viral dsRNA-binding proteins of vaccinia virus (E3L) (33), influenza A virus (NS1) (34), and likely group C rotavirus dsRNA-binding protein (30). Instead, the L* protein expressed by Theiler’s virus and the L^pro^ protein of foot-and-mouth disease virus act by binding to RNase L, preventing its interaction with 2-5A and subsequent dimerization (35, 36). Similar to VP3 PDE, the ns2 protein of murine coronavirus virus and the NS4b protein of Middle East respiratory syndrome coronavirus (MERS-CoV) are phosphodiesterases that hydrolyze 2-5A (13, 37–39). Finally, group C enteroviruses have a phylogenetically conserved RNA structure which functions as a competitive inhibitor of the endoribonuclease domain of RNase L (40). An important advantage of the RNase L degradation mechanism used by SA11 virus is that it does not require stoichiometric expression of NSP1. Operating through its CRL-like activity, even low amounts of NSP1 can move from one RNase L molecule, depleting the entire pool.

## MATERIALS AND METHODS

### Cells and viruses

Baby hamster kidney cells expressing T7 RNA polymerase (BHK-T7) were a kind gift of Dr. Ulla Buchholz (NIAID, NIH). BHK-T7 cells were cultured in Glasgow medium as described previously (41). MA104 and HEK 293T cells were cultured in DMEM medium [Dulbecco’s Modified Eagle’s Medium containing 10% FBS and 1% penicillin-streptomycin mixture]. Incomplete DMEM is the same as complete DMEM, except lacking FBS. HT-29 cells were cultured in McCoy’s 5A medium (HyClone) supplemented with 10% FBS and 1% penicillin-streptomycin mixture.

Rotavirus strains SA11-L2, RRV, and OSU were provided by NIAID, NIH. The monoreassortant rotaviruses SRF, SNF, SOF, SDF and SKF were a generous gift of Dr. Nobumichi Kobayashi, Sapporo Medical University (29). Viruses was propagated and titered by plaque assay using MA104 cells (42).

### Recombinant rotaviruses

WT and mutant rSA11 viruses (Table 1) were generated using the reverse genetics system described by Philip *et al* (41). Briefly, BHK-T7 monolayers were transfected with pT7 plasmids (800 ng each) of pT7 expressing the 11 SA11 (+)RNAs, except that 2.4 μg each of pT7/NSP2SA11 and pT7/NSP5SA11 plasmids were used (43). The pT7/NSP1SA11 and pT7/VP3SA11 plasmids were replaced with corresponding modified plasmids as necessary to generate mutant viruses (Table 1). Plasmid mixtures also included 800 ng of pCMV-NP868R. Three days after transfection, BHK-T7 cells were overseeded with MA104 cells. Four days later, infected cell lysates were harvested and amplified on fresh MA104 cell monolayers. After amplification, recombinant viruses were plaque isolated and the sequences of their modified genome segments confirmed by cDNA sequencing. Recombinant viruses were amplified on MA104 cells, purified by pelleting through a 1 ml cushion of 35% sucrose (w/v) in Tris-buffered saline, pH 7.4 (TBS) at 100,000 × *g* for 2.5 h (4°C) using a Beckman SW28 rotor, and resuspended in TBS. Genomic dsRNAs were recovered from viruses by Trizol extraction, resolved by electrophoresis on 10% polyacrylamide gels, and detected by staining with ethidium bromide.

**Table 1.**
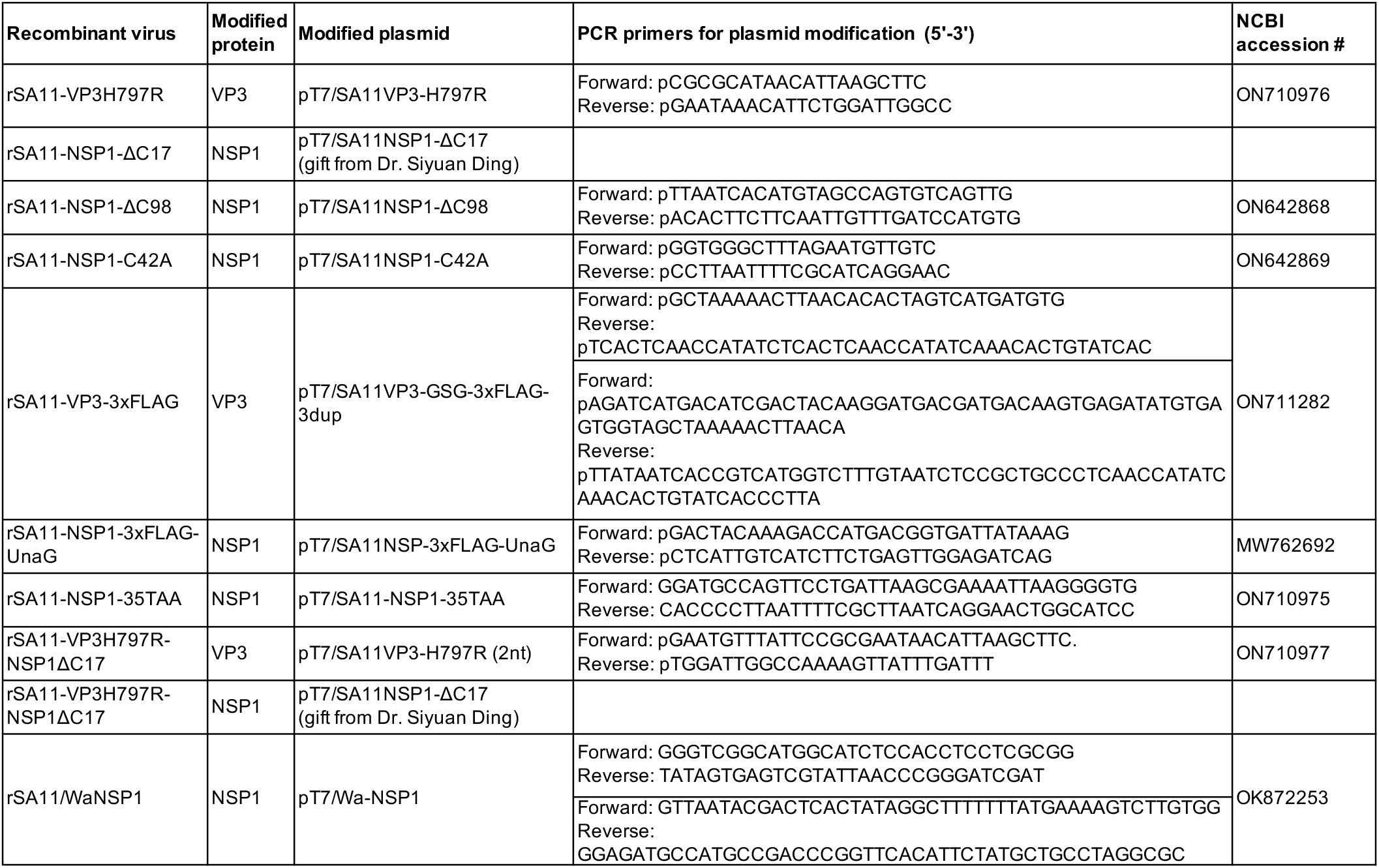
Modified plasmids and primers used in constructing mutant SA11 rotaviruses.

### Multi-step growth curves

Multi-step growth curves of recombinant viruses were performed on MA104 cells in 12-well culture plates. MA104 cells were washed with incomplete DMEM, then infected with virus at a multiplicity of infection (MOI) of 0.01 plaque-forming units (PFU) per ml. After a one-hour adsorption, the cells were washed with incomplete DMEM, overlaid with 1 ml of incomplete DMEM containing 0.5 μg per ml trypsin and incubated until time of harvest. After three rounds of freeze-thaw, infected cell lysates were clarified by centrifugation for 10 min at 500 × *g* (4°C). Virus titers in lysates were determined by plaque assay.

### Plasmids

pT7 plasmids expressing rotavirus SA11 (+)RNAs (pT7/VP1SA11, pT7/VP2SA11, pT7/VP3SA11, pT7/VP4SA11, pT7/VP6SA11, pT7/VP7SA11, pT7/NSP1SA11, pT7/NSP2SA11, pT7/NSP3SA11, pT7/NSP4SA11, and pT7/NSP5SA11) were kindly provided by Dr. Takeshi Kobayashi (Osaka University) {https://www.addgene.org/Takeshi_Kobayashi/} (44). The CMV plasmid expressing the NP868R capping enzyme (pCMV-NP868R) was generated as described previously (45). To generate VP3 PDE and NSP1 mutant viruses, pT7/VP3SA11 and pT7/NSP1SA11 plasmids were modified using a Phusion site-directed mutagenesis kit (ThermoFisher Scientific) and the primers indicated in Table 1. The plasmid pT7/SA11NSP1-ΔC17 was kindly provided by Dr. Siyuan Ding (Washington University in St. Louis). CMV expression vectors used for transient expression of FLAG-tagged human RNase L (pFLAG-CMV2-RNase L), and FLAG-tagged OAS1, OAS2, and OAS3 (p3×FLAG CMV-OAS1 - OAS3, respectively), were kindly provided by Drs. Christina Gaughan and Robert Silverman (Cleveland Clinic) (8, 46). pHTN-HaloTag expression vectors (Promega) encoding different forms of Halo-tagged NSP1 were prepared with a Phusion site-directed mutagenesis kit and the primers and PCR templates indicated in Supplemental Table 1.

### Transfection

To transfect HEK 293T cells, cells were seeded in complete DMEM into 12-well plates. The next day, cells were transfected with 1 μg of a pHTN plasmid expressing WT or mutant NSP1 and 0.3 μg of a pCMV plasmid expressing human RNase L (pFLAG-CMV2- RNase L) using Lipofectamine 2000. The cells were harvested for immunoblot assay at 24 h post transfection.

To transfect HT-29 cells, cells were seeded in complete McCoy’s 5A medium in 12-well plates. The next day, cells were washed with incomplete McCoy’s 5A medium and transfected with 2 μg of a p3×FLAG-CMV plasmid expressing OAS1, OAS2 or OAS3. Six hours later, cells were supplemented with 1 ml of complete McCoy’s 5A medium containing 20% FBS. The day after, cells were infected with WT virus at an MOI of 5 and harvested at 8 h p.i.

### RNA TapeStation assay

MA104 cell monolayers were prepared in 24-well plates using complete DMEM. The next day, the cells were mock infected or infected with rSA11 viruses (MOI=6) using incomplete DMEM. In some cases, complete DMEM was supplemented with 50 ng per ml of rhesus monkey recombinant IFN-β (SinoBiological, 90104-C05H) beginning 24 h prior to infection. In other cases, infected cells were transfected with 500 ng of dsRNA per well at 5 h p.i. At 10 h p.i., infected cells were harvested and total RNA recovered using Trizol. Purified RNA was mixed with RNA ScreenTape sample buffer (Agilent) and analyzed using an Agilent 2200 TapeStation system.

### Inhibitor treatment

One hour before infection, MLN4924 (MilliporeSigma) or MG-132 (Cell Signaling Technology) were added to HT-29 culture media at a final concentration of 1 or 20 μM, respectively. After virus adsorption, cells were washed with serum-free medium and overlaid with the same containing MLN4924 or MG-132 at the concentrations above. Chloroquine (MilliporeSigma) was added to HT-29 culture medium at 3 h p.i. to a final concentration of 50 μM. In experiments with 293T cells, MLN4924 was added to culture medium to a final concentration of 1 μM at 1 h before transfection.

### Immunoblot assay

MA104 or HT-29 cell monolayers were infected with rotaviruses at the indicated MOI. To harvest, the monolayers were washed with cold phosphate-buffered saline, scraped into the same buffer, and pelleted at 5000 × *g* for 10 min (4°C). The pellets were resuspended in lysis buffer [150 mM NaCl, 50 mM Tris-HCl, 1% Triton X-100, 1 × Complete EDTA-free protease inhibitor cocktail (Pierce), pH 7.4]. The lysates were incubated on ice for 30 min, clarified by centrifugation at 15,000 x *g* for 30 min (4°C), mixed with sample buffer (Millipore Sigma) containing SDS and DTT and heated to 95°C for 5 min. Lysate proteins were resolved by electrophoresis on 10%polyacrylamide gels and transferred onto nitrocellulose membranes. The blots were probed with the following primary antibodies: rabbit anti-NSP1 5S (PAC 3977/3978, 1:1000), guinea pig anti-VP6 (Lot 53963, 1:2000), mouse monoclonal anti-Halo (Promega G921A, 1:2000), rabbit monoclonal anti-IFIT1 (CST 14769S, 1:1500), rabbit anti-Viperin (ENZO ALX-210-956-C100, 1:1000), mouse monoclonal anti-RNase L (47)(1:1000), rabbit monoclonal anti-IRF3 (CST 11904S, 1:2000), rabbit monoclonal anti-FLAG (CST 14793S, 1:2000), rabbit monoclonal anti-PCNA (CST 13110S, 1:1000), mouse monoclonal FLAG M2 (Sigma-Aldrich F1804, 1:2000). Primary antibodies were detected using 1:10,000 dilutions of with horseradish peroxidase (HRP)-conjugated secondary antibodies: goat anti-mouse IgG (KPL), goat anti-guinea pig IgG (KPL), or goat anti-rabbit IgG (KPL). Signals were developed using Clarity Western ECL substrate (Bio-Rad 170-5060) and detected using a Bio-Rad ChemiDoc MP imaging system.

### Immunoprecipitation

HEK 293T monolayers in 10-cm dishes were overlaid with complete DMEM containing 1 μM MLN4924. One hour later, the cells were transfected with 2.4 μg pFLAG-CMV2-RNase L and 8 μg pHTN-SA11NSP1 using Lipofectamine 2000. Clarified lysates prepared from the cells at 24 h post transfection were incubated with anti-FLAG M2 magnetic beads (Sigma-Aldrich) or HaloLink resin (Promega). The anti-FLAG beads were washed three times with wash buffer (50 mM Tris [pH 7.4], 150 mM NaCl, 0.1% Tween-20) and divided into two tubes. Beads in one of the tubes were treated with RNase cocktail (Invitrogen) for 15 min at 37°C. Beads were washed once with TBS, and bound proteins were eluted with elution buffer (Pierce). Halo resin was washed three times with wash buffer (50 mM Tris [pH 7.4], 150 mM NaCl, 0.5% Triton X-100 and 1 mg/ml BSA) and once with TBS. The resin was divided into two tubes. Resin in one of the tubes was treated with RNase cocktail, as above. Afterwards, the Halo resins were washed once with TBS and eluted with SDS sample buffer containing DTT. Eluted proteins were resolved by electrophoresis on 10% polyacrylamide gels and detected by immunoblot assay, as described above.

### Mass spectrometry analysis

For one hour prior to infection, HT-29 cells were maintained in complete DMEM containing 20 μM MG-132. Cells were washed three times with incomplete DMEM and infected with trypsin-activated rSA11-VP3-3xFLAG or rSA11-NSP1-3×FLAG-UnaG virus at an MOI of 5. One hour later, cells were washed and overlaid with incomplete DMEM supplemented with MG-132. At 7 h p.i., cells were harvested and clarified lysates prepared as above. Immunoprecipitation using mouse anti-FLAG M2 beads was used to recover complexes from the clarified lysates containing 3×FLAG-tagged VP3 or NSP1. Anti-FLAG M2 magnetic beads containing 3×FLAG-tagged VP3 or 3x-FLAG-tagged NSP1 were sent to the IU Laboratory for Biological Mass Spectrometry and analyzed by LC-MS on an Orbitrap Fusion Lumos equipped with an Easy NanoLC1200. Data were analyzed using Proteome Discoverer (2.5). The data output will be supplied on request.

## ACKNOWLEDGEMENTS

Our appreciation goes out the virologists in the Patton, Danthi, Hardy and Mukhopadhyay laboratories for their support and encouragement. Special thanks to Ashley Long for generating the rSA11/WaNSP1 virus. We acknowledge Dr. Jonathan Trinidad and the IU Laboratory for Biological Mass Spectrometry for help in our mass spectrometry experiments. This work was supported by the National Institutes of Health (R21AI144881, R21AI14271), Indiana Clinical and Translational Sciences Institute, and the Lawrence M. Blatt Endowment.

---

**Supplemental Table 1.**
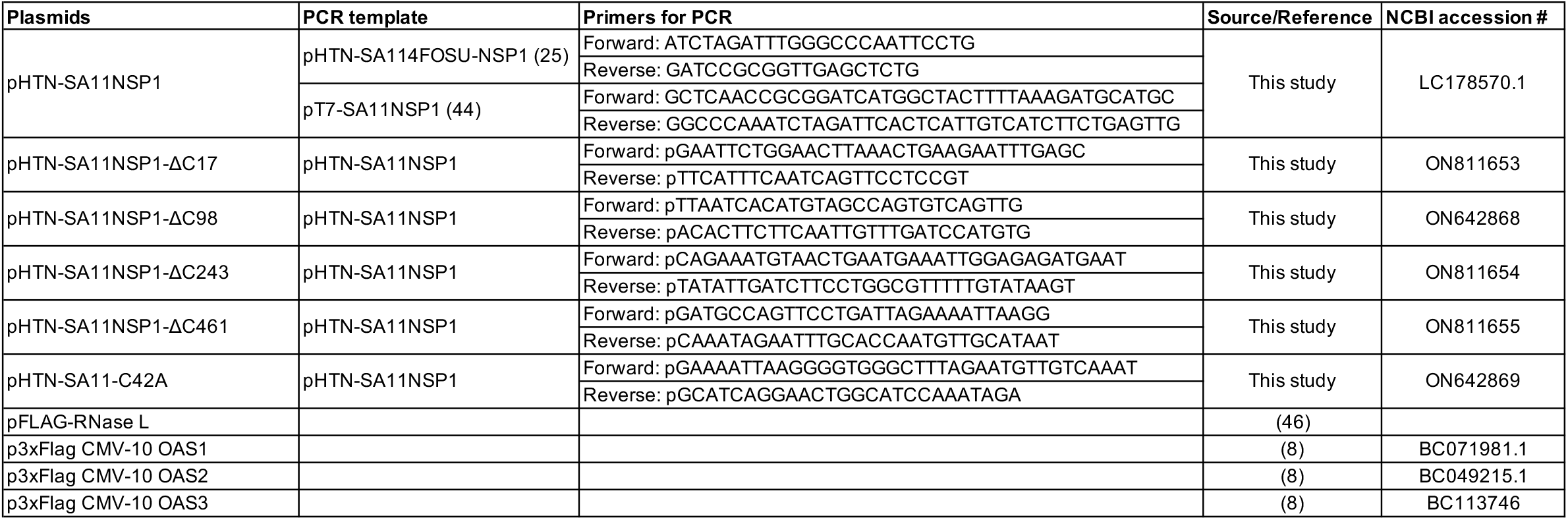
Plasmids used in transient expression of NSP1, RNase L and OAS proteins.

**Figure S1.**
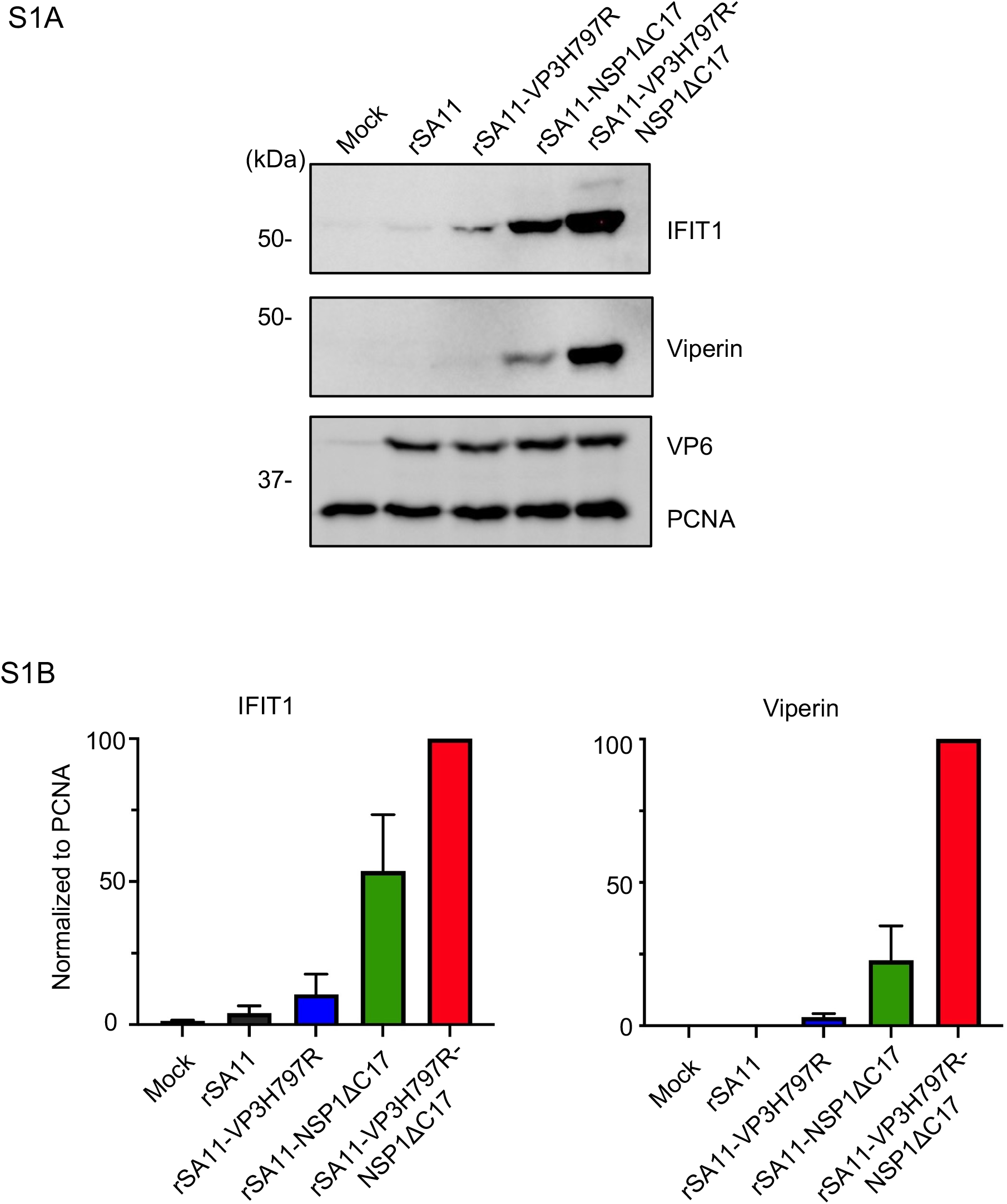
Induction of ISGs in cells infected rotaviruses with mutations in VP3 PDE and NSP1. (A) MA104 cells were infected with the indicated rotavirus strains (MOI = 6). Cells were harvested at 8 h p.i. and analyzed by immunoblot assay for IFIT1, viperin, VP6, and PCNA. (B) Levels of IFIT1 and viperin in blots were determined using ImageJ software. The data are representative of two independent experiments.

## Notes

### Competing Interest Statement

The authors have declared no competing interest.

## REFERENCES

1. Troeger C, Khalil IA, Rao PC, Cao S, Blacker BF, Ahmed T, Armah G, Bines JE, Brewer TG, Colombara DV, Kang G, Kirkpatrick BD, Kirkwood CD, Mwenda JM, Parashar UD, Petri WA, Jr., Riddle MS, Steele AD, Thompson RL, Walson JL, Sanders JW, Mokdad AH, Murray CJL, Hay SI, Reiner RC, Jr. 2018. Rotavirus Vaccination and the Global Burden of Rotavirus Diarrhea Among Children Younger Than 5 Years. JAMA Pediatr 172:958–965.

2. Desselberger U. 2014. Rotaviruses. Virus Res 190:75–96.

3. Trask SD, McDonald SM, Patton JT. 2012. Structural insights into the coupling of virion assembly and rotavirus replication. Nat Rev Microbiol 10:165–77.

4. Zhu S, Ding S, Wang P, Wei Z, Pan W, Palm NW, Yang Y, Yu H, Li HB, Wang G, Lei X, de Zoete MR, Zhao J, Zheng Y, Chen H, Zhao Y, Jurado KA, Feng N, Shan L, Kluger Y, Lu J, Abraham C, Fikrig E, Greenberg HB, Flavell RA. 2017. Nlrp9b inflammasome restricts rotavirus infection in intestinal epithelial cells. Nature 546:667–670.

5. Sen A, Pruijssers AJ, Dermody TS, Garcia-Sastre A, Greenberg HB. 2011. The early interferon response to rotavirus is regulated by PKR and depends on MAVS/IPS-1, RIG-I, MDA-5, and IRF3. J Virol 85:3717–32.

6. Broquet AH, Hirata Y, McAllister CS, Kagnoff MF. 2011. RIG-I/MDA5/MAVS are required to signal a protective IFN response in rotavirus-infected intestinal epithelium. J Immunol 186:1618–26.

7. Schoggins JW, Wilson SJ, Panis M, Murphy MY, Jones CT, Bieniasz P, Rice CM. 2011. A diverse range of gene products are effectors of the type I interferon antiviral response. Nature 472:481–5.

8. Li Y, Banerjee S, Wang Y, Goldstein SA, Dong B, Gaughan C, Silverman RH, Weiss SR. 2016. Activation of RNase L is dependent on OAS3 expression during infection with diverse human viruses. Proc Natl Acad Sci U S A 113:2241–6.

9. Malathi K, Dong B, Gale M, Jr., Silverman RH. 2007. Small self-RNA generated by RNase L amplifies antiviral innate immunity. Nature 448:816–9.

10. Zhou A, Paranjape J, Brown TL, Nie H, Naik S, Dong B, Chang A, Trapp B, Fairchild R, Colmenares C, Silverman RH. 1997. Interferon action and apoptosis are defective in mice devoid of 2′,5′-oligoadenylate-dependent RNase L. EMBO J 16:6355–63.

11. Ogden KM, Snyder MJ, Dennis AF, Patton JT. 2014. Predicted structure and domain organization of rotavirus capping enzyme and innate immune antagonist VP3. J Virol 88:9072–85.

12. Ogden KM, Hu L, Jha BK, Sankaran B, Weiss SR, Silverman RH, Patton JT, Prasad BV. 2015. Structural basis for 2′-5′-oligoadenylate binding and enzyme activity of a viral RNase L antagonist. J Virol 89:6633–45.

13. Asthana A, Gaughan C, Dong B, Weiss SR, Silverman RH. 2021. Specificity and Mechanism of Coronavirus, Rotavirus, and Mammalian Two-Histidine Phosphoesterases That Antagonize Antiviral Innate Immunity. mBio 12:e0178121.

14. Arnold MM, Patton JT. 2011. Diversity of interferon antagonist activities mediated by NSP1 proteins of different rotavirus strains. J Virol 85:1970–9.

15. Lutz LM, Pace CR, Arnold MM. 2016. Rotavirus NSP1 Associates with Components of the Cullin RING Ligase Family of E3 Ubiquitin Ligases. J Virol 90:6036–48.

16. Graff JW, Ewen J, Ettayebi K, Hardy ME. 2007. Zinc-binding domain of rotavirus NSP1 is required for proteasome-dependent degradation of IRF3 and autoregulatory NSP1 stability. J Gen Virol 88:613–620.

17. Barro M, Patton JT. 2005. Rotavirus nonstructural protein 1 subverts innate immune response by inducing degradation of IFN regulatory factor 3. Proc Natl Acad Sci U S A 102:4114–9.

18. Barro M, Patton JT. 2007. Rotavirus NSP1 inhibits expression of type I interferon by antagonizing the function of interferon regulatory factors IRF3, IRF5, and IRF7. J Virol 81:4473–81.

19. Graff JW, Ettayebi K, Hardy ME. 2009. Rotavirus NSP1 inhibits NFkappaB activation by inducing proteasome-dependent degradation of beta-TrCP: a novel mechanism of IFN antagonism. PLoS Pathog 5:e1000280.

20. Ding S, Mooney N, Li B, Kelly MR, Feng N, Loktev AV, Sen A, Patton JT, Jackson PK, Greenberg HB. 2016. Comparative Proteomics Reveals Strain-Specific beta-TrCP Degradation via Rotavirus NSP1 Hijacking a Host Cullin-3-Rbx1 Complex. PLoS Pathog 12:e1005929.

21. Davis KA, Morelli M, Patton JT. 2017. Rotavirus NSP1 Requires Casein Kinase II-Mediated Phosphorylation for Hijacking of Cullin-RING Ligases. mBio 8(4):e01213–17.

22. Morelli M, Ogden KM, Patton JT. 2015. Silencing the alarms: Innate immune antagonism by rotavirus NSP1 and VP3. Virology 479-480:75–84.

23. Bhowmick R, Mukherjee A, Patra U, Chawla-Sarkar M. 2015. Rotavirus disrupts cytoplasmic P bodies during infection. Virus Res 210:344–54.

24. Arnold MM. 2016. The Rotavirus Interferon Antagonist NSP1: Many Targets, Many Questions. J Virol 90:5212–5215.

25. Morelli M, Dennis AF, Patton JT. 2015. Putative E3 ubiquitin ligase of human rotavirus inhibits NF-kappaB activation by using molecular mimicry to target beta-TrCP. mBio 6(1):e02490–14.

26. Sanchez-Tacuba L, Rojas M, Arias CF, Lopez S. 2015. Rotavirus Controls Activation of the 2′-5′-Oligoadenylate Synthetase/RNase L Pathway Using at Least Two Distinct Mechanisms. J Virol 89:12145–53.

27. Song Y, Feng N, Sanchez-Tacuba L, Yasukawa LL, Ren L, Silverman RH, Ding S, Greenberg HB. 2020. Reverse Genetics Reveals a Role of Rotavirus VP3 Phosphodiesterase Activity in Inhibiting RNase L Signaling and Contributing to Intestinal Viral Replication In Vivo. J Virol 94(9):e01952–19.

28. Saxena K, Simon LM, Zeng XL, Blutt SE, Crawford SE, Sastri NP, Karandikar UC, Ajami NJ, Zachos NC, Kovbasnjuk O, Donowitz M, Conner ME, Shaw CA, Estes MK. 2017. A paradox of transcriptional and functional innate interferon responses of human intestinal enteroids to enteric virus infection. Proc Natl Acad Sci U S A 114:E570–E579.

29. Mahbub Alam M, Kobayashi N, Ishino M, Naik TN, Taniguchi K. 2006. Analysis of genetic factors related to preferential selection of the NSP1 gene segment observed in mixed infection and multiple passage of rotaviruses. Arch Virol 151:2149–59.

30. Langland JO, Pettiford S, Jiang B, Jacobs BL. 1994. Products of the porcine group C rotavirus NSP3 gene bind specifically to double-stranded RNA and inhibit activation of the interferon-induced protein kinase PKR. J Virol 68:3821–9.

31. Birdwell LD, Zalinger ZB, Li Y, Wright PW, Elliott R, Rose KM, Silverman RH, Weiss SR. 2016. Activation of RNase L by Murine Coronavirus in Myeloid Cells Is Dependent on Basal Oas Gene Expression and Independent of Virus-Induced Interferon. J Virol 90:3160–72.

32. Li Y, Dong B, Wei Z, Silverman RH, Weiss SR. 2019. Activation of RNase L in Egyptian Rousette Bat-Derived RoNi/7 Cells Is Dependent Primarily on OAS3 and Independent of MAVS Signaling. mBio 94(9):e01952–19.

33. Xiang Y, Condit RC, Vijaysri S, Jacobs B, Williams BR, Silverman RH. 2002. Blockade of interferon induction and action by the E3L double-stranded RNA binding proteins of vaccinia virus. J Virol 76:5251–9.

34. Min JY, Krug RM. 2006. The primary function of RNA binding by the influenza A virus NS1 protein in infected cells: Inhibiting the 2′-5′ oligo (A) synthetase/RNase L pathway. Proc Natl Acad Sci U S A 103:7100–5.

35. Drappier M, Jha BK, Stone S, Elliott R, Zhang R, Vertommen D, Weiss SR, Silverman RH, Michiels T. 2018. A novel mechanism of RNase L inhibition: Theiler’s virus L* protein prevents 2-5A from binding to RNase L. PLoS Pathog 14:e1006989.

36. Sui C, Jiang D, Wu X, Liu S, Li F, Pan L, Cong X, Li J, Yoo D, Rock DL, Miller LC, Lee C, Du Y, Qi J. 2021. Inhibition of Antiviral Innate Immunity by Foot-and-Mouth Disease Virus L(pro) through Interaction with the N-Terminal Domain of Swine RNase L. J Virol 95:e0036121.

37. Zhao L, Jha BK, Wu A, Elliott R, Ziebuhr J, Gorbalenya AE, Silverman RH, Weiss SR. 2012. Antagonism of the interferon-induced OAS-RNase L pathway by murine coronavirus ns2 protein is required for virus replication and liver pathology. Cell Host Microbe 11:607–16.

38. Zhang R, Jha BK, Ogden KM, Dong B, Zhao L, Elliott R, Patton JT, Silverman RH, Weiss SR. 2013. Homologous 2′,5′-phosphodiesterases from disparate RNA viruses antagonize antiviral innate immunity. Proc Natl Acad Sci U S A 110:13114–9.

39. Thornbrough JM, Jha BK, Yount B, Goldstein SA, Li Y, Elliott R, Sims AC, Baric RS, Silverman RH, Weiss SR. 2016. Middle East Respiratory Syndrome Coronavirus NS4b Protein Inhibits Host RNase L Activation. mBio 7:e00258.

40. Townsend HL, Jha BK, Han JQ, Maluf NK, Silverman RH, Barton DJ. 2008. A viral RNA competitively inhibits the antiviral endoribonuclease domain of RNase L. RNA 14:1026–36.

41. Philip AA, Dai J, Katen SP, Patton JT. 2020. Simplified Reverse Genetics Method to Recover Recombinant Rotaviruses Expressing Reporter Proteins. J Vis Exp doi:10.3791/61039.

42. Arnold M, Patton JT, McDonald SM. 2009. Culturing, storage, and quantification of rotaviruses. Curr Protoc Microbiol Chapter 15:Unit 15C 3.

43. Komoto S, Fukuda S, Kugita M, Hatazawa R, Koyama C, Katayama K, Murata T, Taniguchi K. 2019. Generation of Infectious Recombinant Human Rotaviruses from Just 11 Cloned cDNAs Encoding the Rotavirus Genome. J Virol 93(8):e02207–18.

44. Kanai Y, Komoto S, Kawagishi T, Nouda R, Nagasawa N, Onishi M, Matsuura Y, Taniguchi K, Kobayashi T. 2017. Entirely plasmid-based reverse genetics system for rotaviruses. Proc Natl Acad Sci U S A 114:2349–2354.

45. Philip AA, Perry JL, Eaton HE, Shmulevitz M, Hyser JM, Patton JT. 2019. Generation of Recombinant Rotavirus Expressing NSP3-UnaG Fusion Protein by a Simplified Reverse Genetics System. J Virol 93(8):e02207–18.

46. Zhang A, Dong B, Doucet AJ, Moldovan JB, Moran JV, Silverman RH. 2014. RNase L restricts the mobility of engineered retrotransposons in cultured human cells. Nucleic Acids Res 42:3803–20.

47. Dong B, Silverman RH. 1995. 2-5A-dependent RNase molecules dimerize during activation by 2-5A. J Biol Chem 270:4133–7.

